# SARS-CoV-2 spike protein induces inflammation via TLR2-dependent activation of the NF-κB pathway

**DOI:** 10.1101/2021.03.16.435700

**Authors:** Shahanshah Khan, Mahnoush S. Shafiei, Christopher Longoria, John Schoggins, Rashmin C. Savani, Hasan Zaki

## Abstract

Pathogenesis of COVID-19 is associated with a hyperinflammatory response; however, the precise mechanism of SARS-CoV-2-induced inflammation is poorly understood. Here we investigated direct inflammatory functions of major structural proteins of SARS-CoV-2. We observed that spike (S) protein potently induces inflammatory cytokines and chemokines including IL-6, IL-1ß, TNFa, CXCL1, CXCL2, and CCL2, but not IFNs in human and mouse macrophages. No such inflammatory response was observed in response to membrane (M), envelope (E), and neucleocapsid (N) proteins. When stimulated with extracellular S protein, human lung epithelial cells A549 also produce inflammatory cytokines and chemokines. Interestingly, epithelial cells expressing S protein intracellularly are non-inflammatory, but elicit an inflammatory response in macrophages when co-cultured. Biochemical studies revealed that S protein triggers inflammation via activation of the NF-κB pathway in a MyD88-dependent manner. Further, such an activation of the NF-κB pathway is abrogated in Tlr2-deficient macrophages. Consistently, administration of S protein induces IL-6, TNF-a, and IL-1 ß in wild-type, but not Tlr2-deficient mice. Together these data reveal a mechanism for the cytokine storm during SARS-CoV-2 infection and suggest that TLR2 could be a potential therapeutic target for COVID-19.

## INTRODUCTION

Coronavirus induced disease (COVID) 19, caused by severe acute respiratory syndrome coronavirus 2 (SARS-CoV-2), has emerged as a major public health crisis since December 2019 (1–3). SARS-CoV-2 is a positive sense single stranded RNA virus. Like other coronaviruses, such as SARS (retrospectively named SARS-CoV-1) and Middle Eastern respiratory syndrome (MERS)-CoV, SARS-CoV-2 primarily causes infection in the respiratory tract, leading to either asymptomatic infection or a range of symptoms including cough, fever, pneumonia, respiratory failure, along with other complications like diarrhea and multi-organ failure (4–7). In the absence of effective therapies, COVID-19 has caused 1.8 million deaths worldwide by the end of 2020. Although our knowledge is still evolving, immunopathology caused by cytokine storm plays a decisive role in COVID-19 pathogenesis (4, 5, 8).

SARS-CoV-2 infects human cells through its Spike (S) protein, which binds to the receptor angiotensin converting enzyme 2 (ACE2), expressed on alveolar epithelial cells, allowing endocytosis of the viral particle (1, 9). Following endocytosis, the viral genome is replicated using both viral and host machineries, leading to the death of virally infected cells (10). The pathology of SARS-CoV-2 infected lung is further worsened with inflammatory responses of innate immune cells such as macrophages, monocytes, and neutrophils, which are activated by viral components and products of apoptotic and necrotic cells (3). While innate immune response is essential for anti-viral host defense, excessive inflammatory cytokines and chemokines are cytotoxic for respiratory epithelial cells and vascular endothelial cells (11). Indeed, clinical manifestation of COVID-19 is marked by higher levels of IL-2, IL-6, IL-8, TNFa, IFNg, MCP1, MIP1a, IP-10, and GMCSF in patient’s blood (3, 12, 13). Disease severity and death of COVID-19 patients have been correlated to the elevated levels of IL-6 and TNFa (3, 14). However, our understanding of the precise mechanism of induction of proinflammatory cytokines and chemokines during SARS-CoV-2 infection is very limited.

The innate immune inflammatory response is initiated with the recognition of pathogen-associated molecular patterns (PAMPs) by pattern recognition receptors (PRRs), such as Toll-like receptors (TLRs), NOD-like receptors (NLRs), and RIG-I like receptors (RLRs). Activated PRRs involve multiple signaling adapters to activate transcription factors, such as NF-κB, AP1, and IRF3, which regulate the expression of genes involved in immunity and inflammation. RNA sensing receptors such as TLR7, RIG-I, and MDA5 play a central role in anti-viral immunity by inducing type I interferons (IFNa and IFNß) via IRF3 and NF-κB(15–19). Although the relative contribution of RNA sensing pathways in SARS-CoV-2-mediated immunopathology is yet to be explored, previous studies reported that macrophages and dendritic cells infected with SARS-CoV-1 produce proinflammatory cytokines and chemokines, but not type I interferons (20, 21). Consistently, inflammatory responses in severe COVID-19 patients are characterized by high levels of proinflammatory cytokines, but poor type I interferon response (8, 14). Beyond these phenotypic observations, the precise mechanism of the hyperinflammatory response during SARS-CoV-2 infection is poorly understood.

SARS-CoV-2 is an enveloped virus consisting of four major structural proteins – S, nucleocapsid (N), membrane (M), and envelop (E) (22). S protein binds to the receptor binding domain of ACE2 through its S1 subunit allowing proteasomal cleavage of S protein and fusion of the S2 subunit with the host cell membrane (9, 10, 23–25). Thus, SARS-CoV-2 structural proteins are likely to be exposed to PRRs located on the cell membrane, endosome, and cytosol of the infected cell. However, our knowledge on the role of SARS-CoV-2 structural proteins in the innate immune response is very limited. Here, we investigated inflammatory properties of S, M, N and E proteins and revealed that S protein, but not M, N, and E proteins, is a potent viral PAMP, which stimulates macrophages, monocytes, and lung epithelial cells. We demonstrated that S protein is sensed by TLR2, leading to the activation of the NF-κB pathway and induction of inflammatory cytokines and chemokines. This study provides critical insight into molecular mechanism that may contribute to cytokine storm during SARS-CoV-2 infection.

## RESULTS

### SARS-CoV-2 S protein induces inflammatory cytokines and chemokines in macrophages and monocytes

Macrophages play a central role in hyperinflammatory response during SARS-CoV-2 infection(26). To understand whether SARS-CoV-2 structural proteins can activate macrophages and monocytes, we stimulated human monocytic cell THP1-derived macrophages with recombinant S1, S2, M, N and E proteins. Interestingly, both S1 and S2 proteins induced proinflammatory cytokines IL-6, TNFa, and IL-1ß, with S2 being more potent, as measured by real-time PCR and ELISA (**Figure 1A and Figure S1A**). Chemokines produced by macrophages and monocytes recruit T cell and other immune cells in the inflamed tissue, aggravating inflammatory damage(4, 26). Both S1 and S2 subunits of S protein induced chemokines including *CXCL1, CXCL2,* and *CCL2* in THP1 cells (**Figure 1A**). Three other structural proteins - M, N and E - did not induce any cytokines and chemokines (**Figure 1A**). Interferons are critical for adaptive immune response and anti-viral immunity (16). However, THP1 cells did not express either type I (IFNa and IFNß) and Type II (IFNg) interferons in response to any of the SARS-CoV-2 structural proteins (**Figure 1A**). Notably, THP1 cells are not defective in producing interferons when activated by PolyI:C, a ligand for TLR3 (Figure S1B).

**Figure 1.**
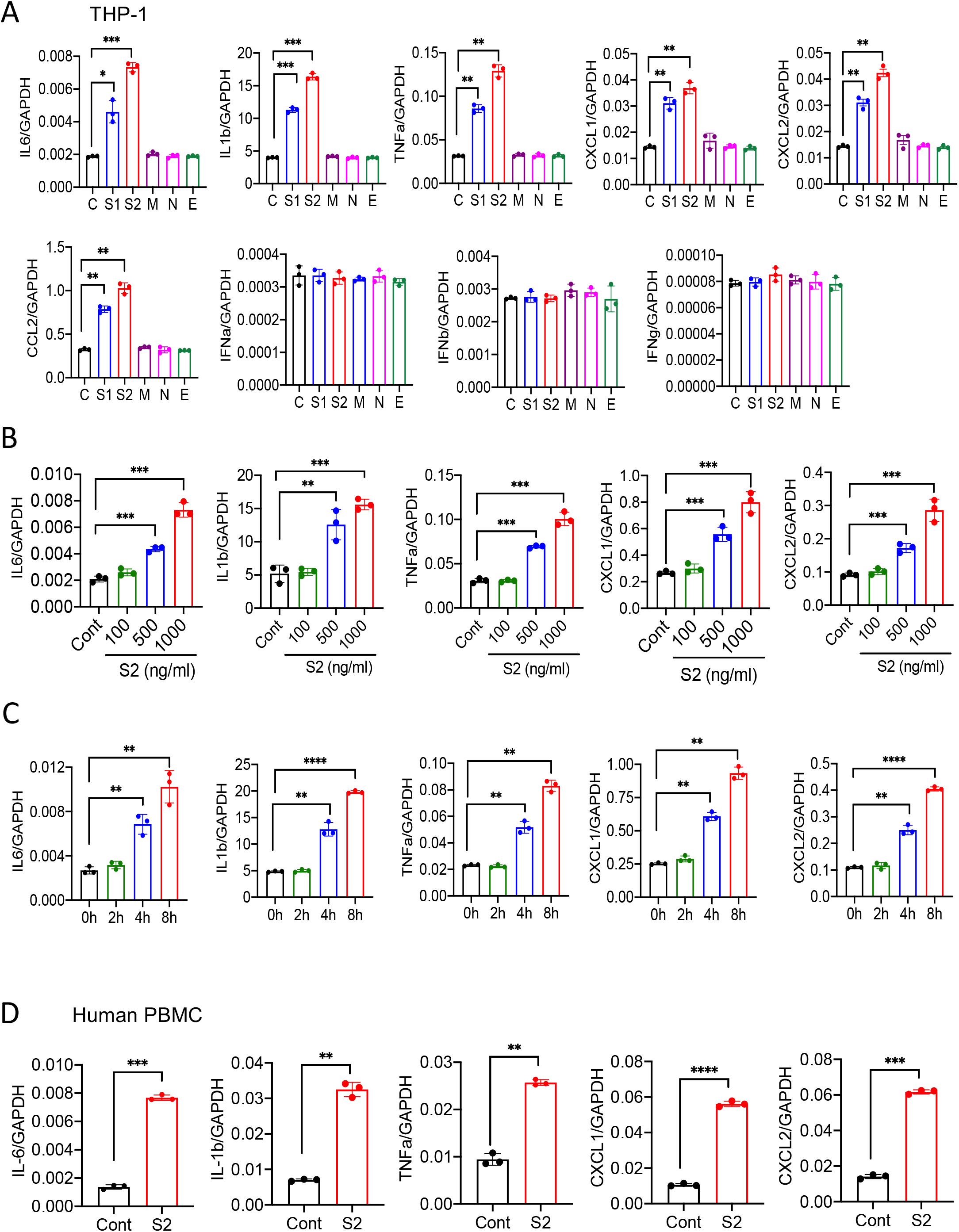
SARS-CoV-2 S protein induces cytokines and chemokines in macrophages and monocytes. (A) Human monocytic cells THP1-derived macrophages were stimulated with recombinant S1, S2, M, N, and E proteins of SARS-CoV-2 at a concentration of 500 ng/ml. Four hours post stimulation, the expression of *IL6, IL1b, TNFa, CXCL1, CXCL2, CCl2, IFNa, IFNb,* and *IFNg* was measured by real-time qPCR. (B) THP-1 cells were stimulated with S2 protein (500ng/ml). At indicated times, RNA was isolated and measured for *IL6, IL1b, TNFa, CXCL1,* and *CXCL2* by real-time qPCR. (C) THP-1 cells were stimulated with S2 protein at various concentration for 4 h, and measured the indicated cytokines by real-time qPCR. (D) Human peripheral blood mononuclear cells (PBMC) were incubated with S2 protein for 4h. The expression of *IL6, IL1b, TNFa, CXCL1,* and *CXCL2* was measured by real-time qPCR. Data represent mean ± SD (n=3); *\*p* < 0.05, \*\**p* < 0.001, *\*\*\*p* < 0.0001, \*\*\*\**p* < 0.00001 by unpaired Student’s *t* test. Experiments were repeated two times and data of a representative experiment is presented.

The expression of cytokines and chemokines in response to S protein was dose dependent (**Figure 1B**). Similarly, S protein-induced inflammatory response was time dependent, with highest being at 8h post stimulation (**Figure 1C**). Notably, heat-denatured S2 protein failed to stimulate THP-1 cells, confirming the specificity of S protein and requirement of its native structural configuration in inducing inflammatory response (Figure S1C). To obtain a direct evidence that S protein can induce inflammatory mediators in human immune cells, we stimulated human peripheral blood mononuclear cells (hPBMC) with S2. There was robust induction of *IL1b, IL6, TNFa, CXCL1,* and *CXCL2* in hPBMC at 4 h post stimulation (**Figure 1D**).

SARS-CoV-2 do not infect mouse cells since S protein cannot bind mouse ACE2 receptor. To understand whether recognition of S protein by ACE2 is required for induction of inflammatory molecules, we stimulated mouse bone marrow-derived macrophages (mBMDMs) with S1 and S2. Interestingly, both S1 and S2 proteins stimulated mBMDMs, expressing *Il6, Il1b, Tnfa, Cxcl1 and Cxcl2* (**Figure S2A**). Similar to THP1 cells, mBMDMs did not produce type I and Type II interferons in response to S protein (**Figure S2A**). Murine macrophage cell line RAW264.7 cells also responded to S2 protein, producing *Il6*, *Tnfa,* and *Il1b* (Figure S2B). Taken together, SARS-CoV-2 S protein potentially induces proinflammatory cytokines and chemokines in macrophages and monocytes.

### Epithelial cells produce inflammatory mediators in response to SARS-CoV-2 S protein

SARS-CoV-2 primarily infect epithelial cells of the lung, kidney, intestine, and vascular endothelial cells (9, 27–29). However, it is poorly understood whether SARS-CoV-2-infected epithelial cells produce proinflammatory cytokines and contribute to cytokine storm of COVID-19 patients. To address this concern, we stimulated human lung cancer epithelial cells A549 and human embryonic kidney epithelial cells HEK293T cells with S1 or S2 proteins. However, there was no induction of *IL6, IL1b, TNFa, CXCL1,* and *CXCL2* in either of the epithelial cells in response to S proteins at 4 h post stimulation (**Figure S3**). Given that epithelial cells are weaker than innate immune cells in expressing inflammatory mediators, we wondered whether the expression of inflammatory molecules in A549 cells is delayed. We, therefore, measured cytokines and chemokines in A549 cells following stimulation with S1 and S2 at 12 and 24h. Interestingly, both S1 and S2 proteins induced proinflammatory cytokines *IL6*, *IL1b, TNFa,* and chemokines *CXCL1* and *CXCL2,* with highest being at 24h post stimulation (**Figure 2A**). IFNγ, but not Type I interferons, was poorly induced by S2 protein in A549 cells (**Figure 2A**).

**Figure 2.**
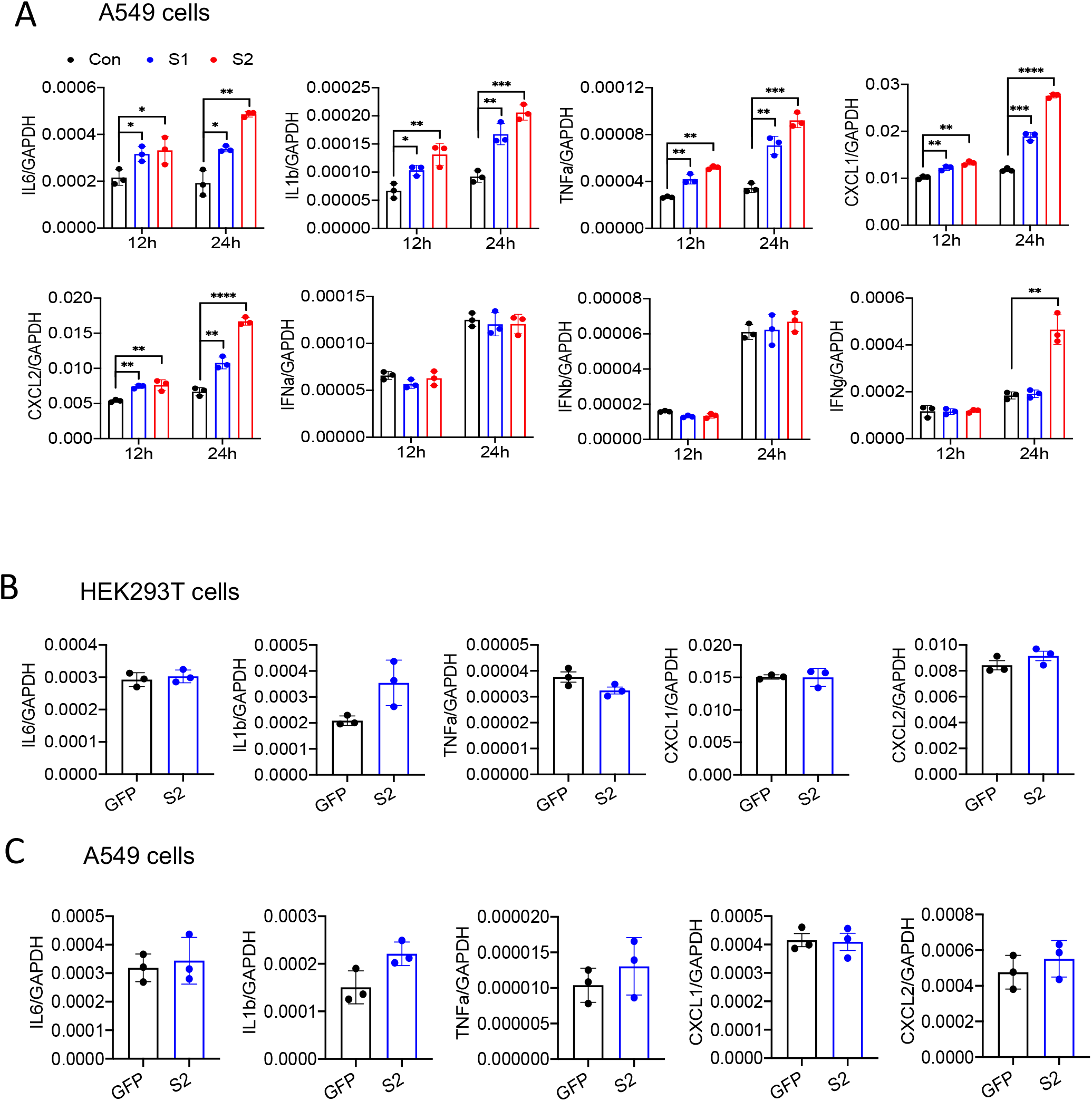
Lung epithelial cells produce inflammatory molecules in response to SARS-CoV-2 S protein. (A) A549 cells were incubated with SARS-CoV-2 S2 (500 ng/ml) protein for 12 and 24h. RNA was isolated and measured for the expression of inflammatory cytokines and chemokines. (B-C) SARS-CoV-2 S protein was overexpressed in HEK293T and A549 cells. 48h following transfection with expression plasmids, the mRNA levels of *IL6, IL1b, TNFa, CXCL1,* and *CXCL2* were measured. Data represent mean ± SD (n=3). Experiments were repeated two times and data of a representative experiment is presented.

We next verified whether cytosolic S protein can stimulate inflammatory response in epithelial cells. Therefore, we transfected HEK293T and A549 cells with plasmids expressing flag-tagged S protein or green florescent protein (GFP). The expression of S protein in HEK293T and A549 cells was confirmed by Western blotting and ELISA (**Figure S4A-C**). However, cytosolic expression of S protein did not induce any cytokines and chemokines in HEK293T cells or A549 cells (**Figure 2B and 2C**). These data suggest that intracellular S protein does not induce inflammatory responses in epithelial cells.

### Epithelial cells expressing S protein trigger inflammation in macrophages

Since airway and other epithelial cells are primary target of SARS-CoV-2, we wondered whether virally infected epithelial cells trigger inflammatory response in macrophages and monocytes in a paracrine manner. To address this concern, we collected culture supernatant of HEK293T-S cells or A549-S cells and added (30% V/V) into the culture medium of THP1 cells (**Figure 3A**). However, culture supernatants of S protein-expressed epithelial cells failed to induce cytokines IL-6, IL-1 ß, and TNFa in THP1 cells (**Figure 3B and 3C**). In fact, S protein was not detectable in the culture supernatant of HEK293T and A549 cells expressing S protein (**Figure S4C**). Flow cytometric analysis suggested that S protein is primarily located in the cytoplasm, not on the cell surface (**Figure S4D**).

**Figure 3.**
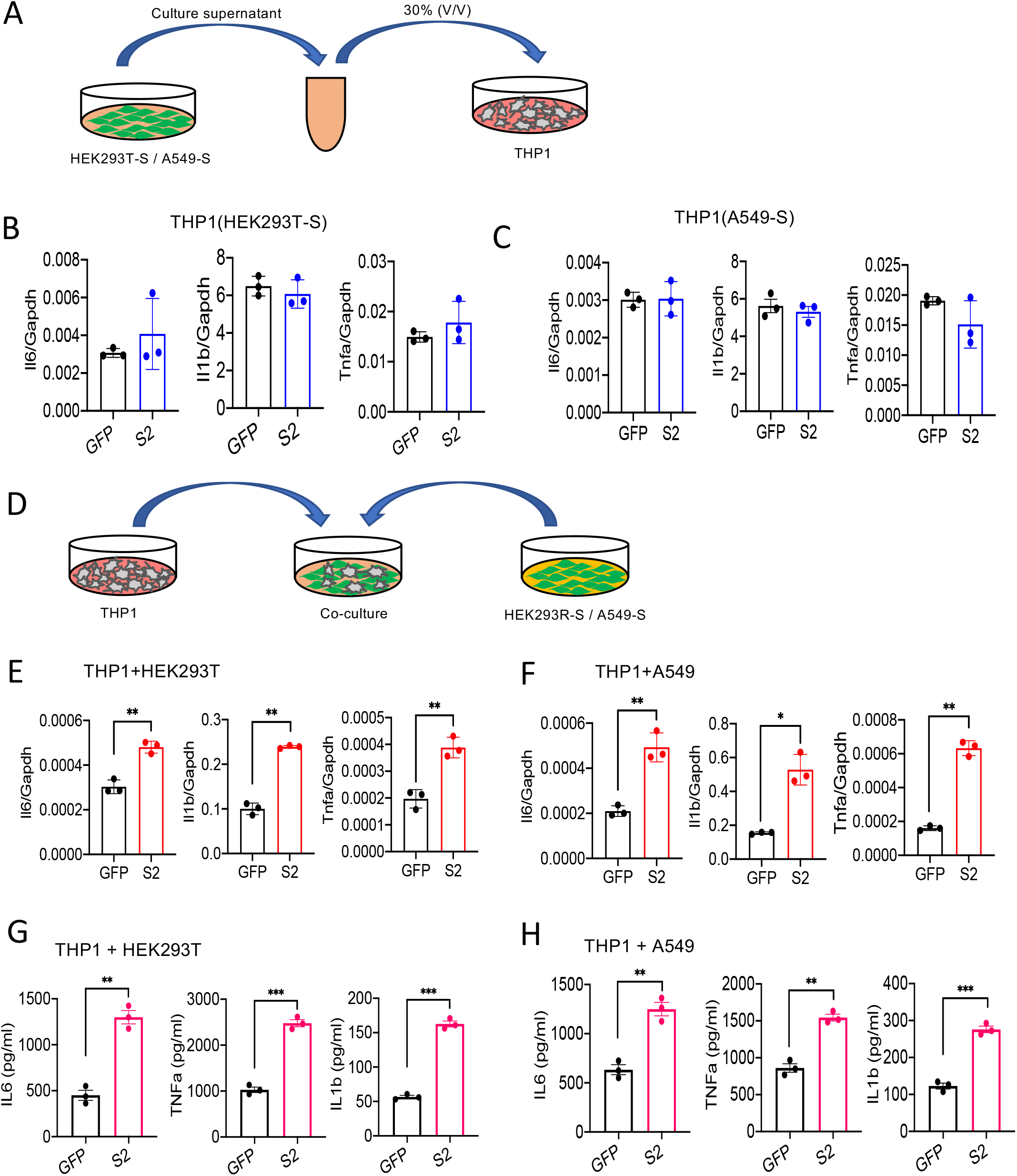
Macrophages get stimulated by epithelial cells expressing S protein. (A-C) SARS-CoV-2 S2 protein was overexpressed in either HEK293T or A549 cells. Forty eight hours following transfection with S2 or GFP plasmids, cell culture supernatants were collected and added into THP1 cell cultures at 30% V/V. (B-C) 4h post incubation, the expression of *IL6, IL1b,* and *TNFa* in THP1 cells treated with culture supernatant of HEK293T (B) or A549 (C) was measured by real-time qPCR. (D) HEK293 or A549 cells expressing S protein were co-cultured with THP1 cells at 1:2 ratio for 16 h. (E-F) The expression of *IL6, IL1b,* and *TNFa* were measured by realtime PCR. (G-H) Protein levels of *IL-6, IL-1ß,* and *TNFa* in culture supernatant described in D were measured by ELISA. Data represent mean ± SD (n=3); \**p* < 0.05, \*\**p* < 0.001, \*\*\**p* < 0.0001 by unpaired Student’s *t* test. Experiments were repeated two times and data of a representative experiment is presented.

We then sought to examine whether innate immune cells become activated when they physically interact with S protein-expressed epithelial cells. Hence, we co-cultured HEK293T-S or A549-S cells with THP1 cells at 1:2 ratio (**Figure 3D)**. Impressively, inflammatory cytokines were appreciably induced in co-cultured cells (**Figure 3E-F**). We confirmed protein levels of IL-6, IL-1ß, and TNFα in the culture supernatant of co-cultured cells by ELISA (**Figure 3G and H**). Notably, similar to Figure S4A-C, S protein was not detected in the culture supernatant of cocultured cells as measured by ELISA (data not shown), suggesting that THP1 cells are activated by S protein-transfected epithelial cells through an unknown mechanism. To further confirm that S protein expressing epithelial cells can stimulate macrophages, we lysed HEK293T-S cells, and added the cell lysate to THP-1cells in culture (**Figure S5A**). Lysates of HEK293T-S cells efficiently induced inflammatory cytokines in THP-1 cells while no such induction was observed in response to cell lysate of HEK293T-GFP (**Figure S5B**). Together, these data imply that SARS-CoV-2 infected epithelial cells may stimulate macrophages and monocytes in a paracrine manner to produce inflammatory mediators.

### S protein activates the NF-κB pathway

Inflammatory genes are transcriptionally regulated by transcription factors that are activated by signaling pathways such as NF-κB, MAPK, STAT3, and AKT. To obtain further insight into how S protein induce the expression of inflammatory mediators, we stimulated THP1 cells with S2 protein. Cell lysates collected at various times following stimulation were analyzed for the activation of these inflammatory pathways by western blotting. As shown in Figure 4A, P65 and IκB were phosphorylated in cells treated with S2 protein. The activation of NF-κB pathway is often accompanied by the activation of MAPK pathways, including ERK, P38 and JNK. Surprisingly, there was no activation of ERK and JNK in S2 stimulated cells (**Figure 4A**). There was no activation of the AKT pathway as well (**Figure 4A**), while STAT3 was phosphorylated at 2h following stimulation (**Figure 4A**). Inflammatory cytokines, such as IL-6, can activate STAT3; thus, the observed activation of STAT3 could be a secondary response of S protein-mediated activation of the NF-κB pathway. S2 protein also activated the NF-κB and STAT3 pathways in A549 cells (**Figure 4B**). To confirm that S protein-induced inflammation was NF-κB dependent, we inhibited the NF-κB pathway using Sc514, an inhibitor of IKKß, during stimulation with S protein. As expected, inhibition of the NF-κB pathway abrogated inflammatory responses in S protein-stimulated macrophages (**Figure 4C and D**).

**Figure 4.**
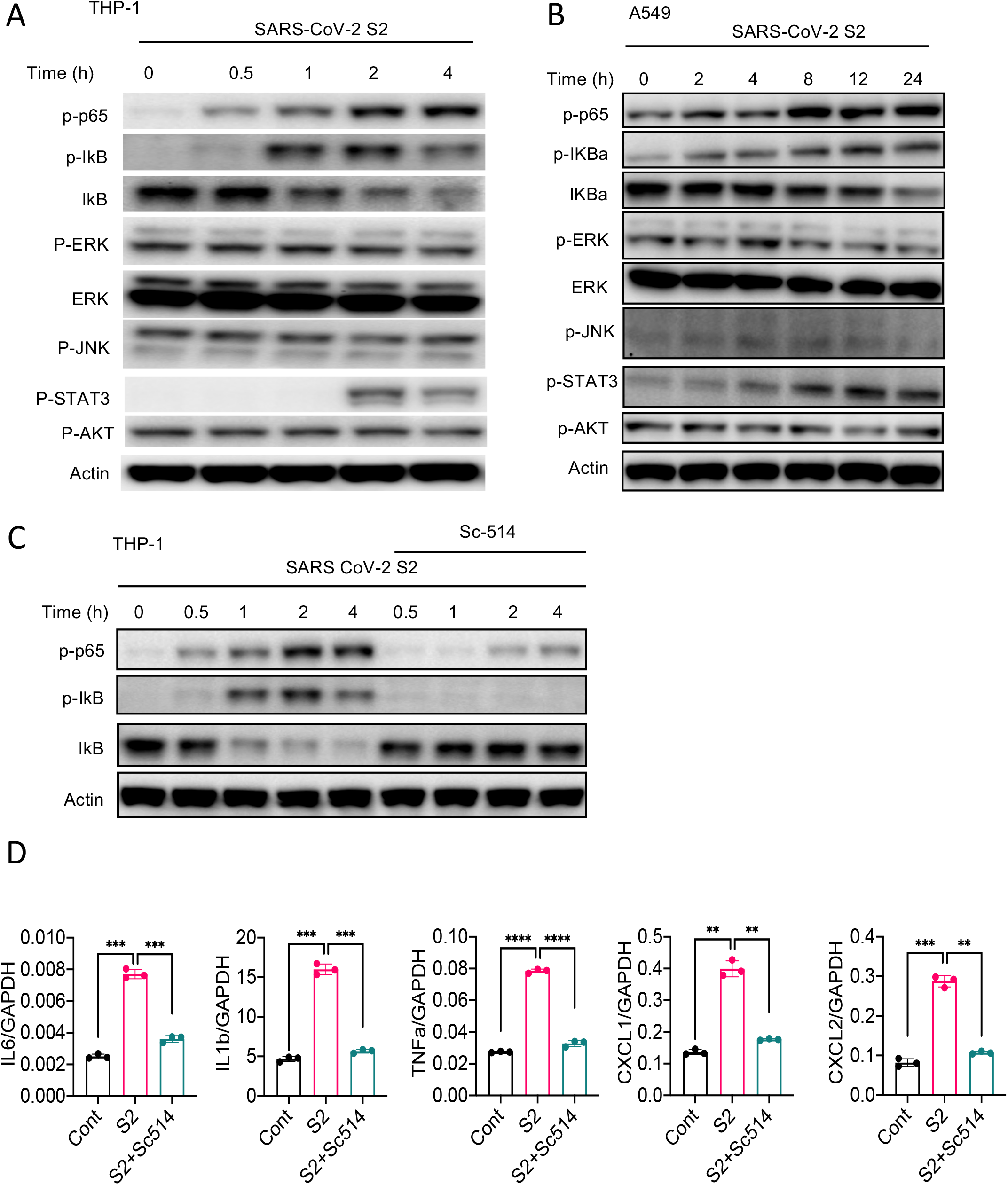
SARS-CoV-2 S protein activates the NF-κB pathway. (A-B) THP1 and A549 cells were stimulated with S2 (500ng/ml) for indicated time points. Phosphorylation of P65, IkBa, ERK, JNK, STAT3, and AKT was measured by western blotting. (C-D) THP1 cells were stimulated by SARS-CoV-2 S2 protein (500ng/ml) in the presence or absence of IKKß inhibitor sc514. (B) Phosphorylation of P65 and IkBa was measured by Western blotting. (E) The expression of *IL6, IL1b* and *TNFa* in stimulated THP1 cells was measured by real-time qPCR. Data represent mean ± SD (n=3); \**p* < 0.05, \*\**p* < 0.001, \*\*\**p* < 0.0001 by unpaired Student’s *t* test. Experiments were repeated two times and data of a representative experiment is presented.

### S protein-mediated activation of the NF-κB pathway is TLR2 dependent

Upon recognition of diverse PAMPs at the cell surface or in the endosome, TLRs activate the NF-κB and MAPK pathways through the adapter protein MyD88. To verify if TLR pathways are involved in S protein-mediated activation of the NF-κB pathway, we stimulated WT and *Myd88^−/−^* BMDM with S2. Interestingly, there was no activation of the NF-κB pathway in *Myd88^−/−^* BMDM (**Figure 5A**). Consistently, there was no cytokine expression in *Myd88^−/−^* macrophages upon stimulation with S protein (**Figure 5B**). This observation suggests that S protein-mediated activation of the NF-κB pathway involves TLR/MyDD88. We then interrogated which TLR sense S protein. Since S is a glycoprotein, we anticipated that TLR2, a receptor for lipoprotein, or TLR4, which senses lipopolysaccharide and several other stimuli (18), could be the immune sensor for S protein. Therefore, we stimulated *Tlr2^−/−^* and *Tlr4^−/−^* BMDMs with S2 protein and measured the activation of the NF-κB pathway. There was no activation of the NF-κB pathway in *Tlr2^−/−^* BMDM, while activation of this pathway was intact in *Tlr4^−/−^* macrophages (**Figure 5C**). We confirmed that *Tlr2^−/−^* macrophages were defective in sensing Pam3CSK4, a ligand for TLR2, while they were fully responsive to TLR4 ligand LPS (**Figure S6**). Similar to S2, S1 protein activated the NF-κB pathway in WT macrophages but not *Tlr2^−/−^* macrophages (**Figure 5D**). Consistent with the defective activation of the NF-κB pathway, there was no induction of proinflammatory cytokines in S2-stimulated *Tlr2^−/−^* macrophages (**Figure 5E**), suggesting that S protein induces inflammatory molecules via TLR2-dependent activation of the NF-κB pathway.

**Figure 5.**
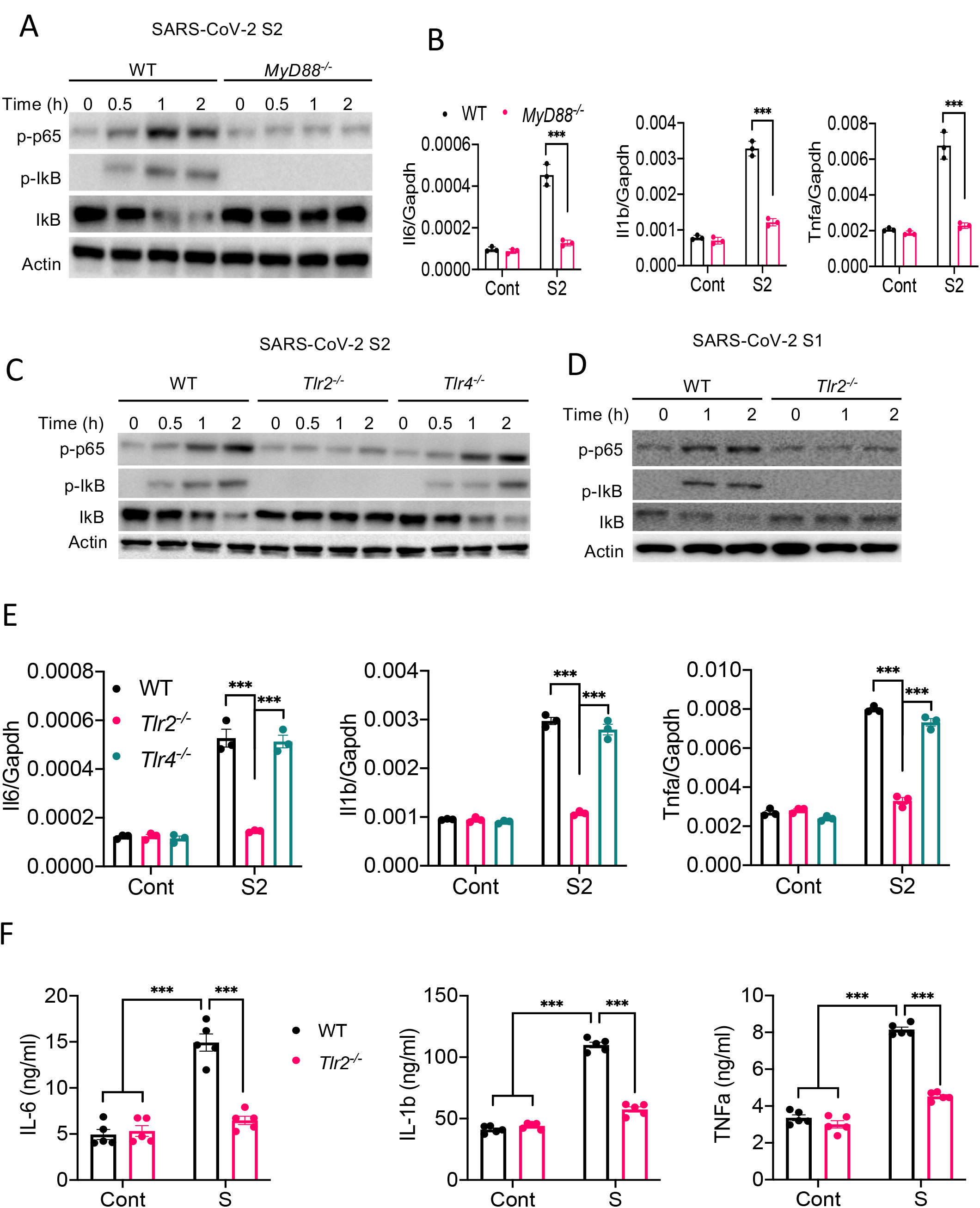
TLR2 recognizes SARS-CoV-2 S protein and activate the NF-κB pathway. (A-B) Bone marrow derived macrophages (BMDMs) from WT and *Myd88^−/−^* mice were stimulated with S2 protein (500 ng/ml). (A) The activation of the NF-κB pathway was measured by western blot analysis of P-P65 and P-Ikba. (B) The induction of *Il6*, *Il1b* and *Tnfa* was measured by real-time PCR. (C) BMDMs from WT, *Tlr2^−/−^* and *Tlr4^−/−^* mice were treated with S2 protein (500 ng/ml). Cell lysates collected at different times were analyzed for the activation of the NF-κB pathway by western blotting of P-P65 and P-IkBa. (D) BMDMs from WT and *Tlr2^−/−^* mice were treated with S1 protein, and the activation of P65 and IkB was measured by western blotting. (E) WT, *Tlr2^−/−^* and *Tlr4^−/−^* macrophages were treated with S2 protein (500ng/ml). The expression of cytokines was measured by real-time PCR at 4 h post stimulation. Data represent mean ± SD (n=3); \*\*\**p* < 0.0001, \*\*\*\**p* < 0.00001 by unpaired Student’s *t* test. Experiments were repeated two times and data of a representative experiment is presented. (F) WT and *Tlr2^−/−^* mice were administered with S1 and S2 protein (1μg each/mouse). Blood collected before and 16h post S protein administration was measured for IL-6, IL-1ß, and TNFa by ELISA. Data represent mean ± SEM (n=5); \*\*\**p* < 0.0001, \*\*\*\**p* < 0.00001 by unpaired Student’s *t* test. Experiments were repeated two times and data of a representative experiment is presented.

Finally, to understand whether S protein induce inflammation in vivo and what role TLR2 plays, we administered S1 and S2 proteins into WT or *Tlr2*^−/−^ mice intraperitoneally (i.p.). 16h following administration of S protein, we measured the cytokine IL-6, IL-1 ß and TNFa in the blood by ELISA before and after challenge with S protein. As shown in **Figure 5F**, the concentrations of IL-6, IL-1ß, and TNFa were elevated following S protein administration in WT mice, whereas no such induction of these cytokines was observed in *Tlr2*^−/−^ mice (**Figure 5F**). Together, these data suggest that TLR2 is the immune sensor for SARS-CoV-2 S protein, which potentially triggers inflammatory responses through the activation of the NF-κB pathway.

## DISCUSSION

Both SARS-CoV-2 infection and aberrant host immune responses are responsible for COVID-19 pathogenesis (4, 5, 8, 26). The initial host immune response against SARS-CoV-2 infection involves innate immune cells, such as macrophages, monocytes, neutrophils, and dendritic cells (30, 31). Cytokines, chemokines, and other inflammatory mediators produced by these cells inhibit virus replication, heal the damage, and activate the adaptive immune system. However, uncontrolled release of cytokines, chemokines, and reactive oxygen and nitrogen species often exert pathological consequences such as tissue injury, systemic inflammation, and organ failure (4, 5, 8). Non-surviving COVID-19 patients exhibited massive influx of macrophages and neutrophils, but reduced T cells in their blood (30), pointing to the association of hyperactivation of innate immune cells with COVID-19 pathogenesis. Indeed, innate immune response is heightened in the lung of COVID-19 patients(31, 32). A better understanding of the mechanism through which SARS-CoV-2 stimulates innate immune cells and activates inflammatory signaling pathways are key to finding better treatment regimens for COVID-19. Our finding that SARS-CoV-2 S protein is a potent viral PAMP involved in the induction of inflammatory cytokines and chemokines via TLR2-dependent activation of the NF-κB pathway, therefore, is a valuable addition to the tremendous scientific effort aiming at combating COVID-19.

Being an RNA virus, SARS-CoV-2 may activate RNA sensors TLR7, RIG-I and MDA5, which primarily responsible for production of type I interferons (17). Interestingly, type I interferon response is attenuated in COVID-19 patients and SARS-CoV-2 infected cells (8, 14). Transcriptomic analysis of bonchoalveolar lavage fluid and peripheral blood mononuclear cells of COVID-19 patients also demonstrated higher expression of proinflammatory cytokines and chemokines, but not type I interferons (32). Our knowledge on type I interferon response of SARS-CoV-2 infected macrophages is limited. However, previous studies on SARS-CoV-1, which share 80% similarity with SARS-CoV-2, showed that macrophages and dendritic cells infected by SARS-CoV-1 produce chemokines CXCL10 and CCL2, but not type I interferons (20, 21). Lack of an interferon response can be explained by the fact that several structural and non-structural proteins including M, N, PLP, ORF3b, ORF6, and NSP1 inhibit type I interferon signaling (33–38). Despite this evidence, the precise mechanism of excessive production of inflammatory cytokines along with reduced of type I interferons in COVID-19 patients remains elusive. In this regard, our findings that SARS-CoV-2 S protein is a potential trigger for proinflammatory cytokines and chemokines help understand why inflammatory response of COVID-19 is marked by elevated levels of proinflammatory cytokines and chemokines, but poor type I interferon response. Our findings suggest that S protein of SARS-CoV-1 and SARS-CoV-2 shares similar inflammatory function (39, 40). Further studies are required to clarify relative contribution of S protein, viral RNA, and other non-structural proteins in COVID-19 associated cytokine storm.

Inflammatory responses of COVID-19 patients are mostly implicated to innate immune cells, but they weakly express ACE2 (41). There is no strong evidence that SARS-CoV-2 infect and propagate in immune cells. Thus, it is intriguing how innate immune cells become activated to produce inflammatory mediators during SARS-CoV-2 infection. We propose three mechanisms involved in hyperinflammatory response during SARS-CoV-2 infection. First, innate immune cells like macrophages and monocytes recognize S protein of SARS-CoV-2 at the cell surface through TLR2, leading to the activation of the NF-κB pathway. Immune sensing of S protein is likely independent of ACE2 since mouse macrophages, whose ACE2 receptor does not bind to S protein, express inflammatory cytokines and chemokines in response to S protein. Second, innate immune cells get activated by virally infected epithelial cells. Our data suggest that epithelial cells expressing S protein in the cytosol can activate macrophage when they physically interact. Although the underlying mechanism is not clear, macrophage may engulf or recognize cell surface molecule expressed on SARS-CoV-2 infected epithelial cells. In a third mechanism, like myeloid cells, epithelial cells can be activated by S protein extracellularly, leading to the induction of proinflammatory cytokines and chemokines. Although inflammatory responses of epithelial cells are weaker than that of innate immune cells, epithelial cell-derived chemokines recruit neutrophils, monocytes and lymphocytes in SARS-CoV2-infected lungs and thereby contribute to immunopathology of COVID-19 patients.

Our data demonstrate that TLR2 is the innate immune sensor for the S protein. Like other MyD88-dependent TLR pathways, ligation of TLR2 leads to the activation of transcription factors NF-κB and AP-1 (17). Interestingly, while there was activation of NF-κB, AP-1 upstream signaling kinases such as ERK and JNK were not seen activated by S proteins. SARS-CoV-1 S protein activates the NF-κB pathway in human monocyte derived macrophages (39, 40). COVID-19 patients also exhibited increased activation of the NF-κB pathway (14). Interestingly, in contrast to these findings, a separate study reported that SARS-CoV-1 S protein expressing baculovirus activates AP-1 but not NF-κB in A549 cells (42). Future studies dissecting the signaling pathway regulated by S protein of SARS-CoV-1 and CoV-2 may reveal further insight.

In summary, this study documents a potential mechanism for the inflammatory response induced by SARS-CoV-2. We demonstrate that SARS-CoV-2 S protein is a potent viral PAMP that upon sensing by TLR2 activates the NF-κB pathway, leading to the expression of inflammatory mediators in innate immune and epithelial cells. The effort so far in combating the COVID-19 pandemic is unprecedented, making it possible for the development of a number of vaccines within a year of outbreak. Since S protein is being targeted by most of the vaccine candidates, it is important to consider its inflammatory function in vaccine design. Considering the fact that new variants of SARS-CoV-2 with mutations in the S protein spread more easily and may confer more severe disease, the effectiveness of current vaccines remain uncertain (43). Thus, the importance of developing therapeutic drugs for COVID-19 remains high. This study suggests that TLR2 or its downstream signaling adapters could be therapeutically targeted to mitigate hyperinflammatory response in COVID-19 patients.

## MATERIALS AND METHODS

### Mice

C57BL6/J (WT), *Myd88^−/−^, Tlr2^−/−^,* and *Tlr4*^−/−^ mice (all C57BL6/J strain), purchased from Jackson Laboratory were used in this study. All mice were bred and maintained in a specific pathogen-free (SPF) facility at the UT Southwestern Medical Center. All studies were approved by the Institutional Animal Care and Use Committee (IACUC) and were conducted in accordance with the IACUC guidelines and the National Institutes of Health Guide for the Care and Use of Laboratory Animals. All experimental groups were conducted with age and sex-matched male and female mice.

### Cell culture and maintenance

The human embryonic kidney epithelial cell line HEK293T (ATCC, CRL-3216), and human lung epithelial cell line A549 (ATCC, CCL-185) were cultured in Dulbecco’s Modified Eagle’s medium (DMEM; high glucose, Sigma) supplemented with 10% (v/v) FBS (Sigma) and 1% (v/v) PenStrep (Sigma) and maintained in a 5% CO2 incubator at 37°C. THP1 (ATCC, CRL-TIB-202) cells were cultured in Roswell Park Memorial Institute (RPMI)-1640 medium (R8758, Sigma) supplemented with 10% (v/v) FBS (Sigma) and 1% (v/v) PenStrep (Sigma) and maintained in a 5% CO2 incubator at 37°C. Each cell lines were confirmed free from mycoplasma contamination by testing with mycoplasma detection kit (Sigma).

### In vitro studies with THP-1 macrophage-like cells

THP-1 cells were cultured in culture medium containing final concentration of 100 ng/mL of phorbol-12-myristate 13-acetate (PMA; tlrl, Invivogen). Following 24h post PMA treatment, THP-1 macrophage-like cells were washed with pre-warmed RPMI-1640 containing 10% FBS and 1% penicillin-streptomycin and allowed to grow in PMA-free culture medium for next 12 hrs. To examine the effect of SARS-CoV-2 structural proteins on inflammatory responses, THP-1 macrophage-like cells were stimulated with SARS-CoV-2 S1 (RayBiotech, 230-30161), SARS-CoV-2 S2 (RayBiotech, 230-30163), SARS-CoV-2 N (RayBiotech, 230-30164), SARS-CoV-2 M (MyBioSource, MBS8574735) and SARS-CoV-2 E (MyBioSource, MBS9141944) for 4 hrs.

### Culture of mouse bone marrow-derived macrophage (BMDM)

Mouse bone marrow cells were collected as described previously (44). Bone marrow cells were cultured in L-cell-conditioned IMDM medium supplemented with 10% FBS, 1% nonessential amino acid, and 1% penicillin-streptomycin for 6 days to differentiate into macrophages. BMDMs were seeded in 12-well cell culture plates at a concentration of 1.2×10^6^ cell/well and incubated overnight before in vitro studies. BMDMs were stimulated as described above.

### cDNA constructs and transient transfection

At 50 - 60 % confluency, HEK293T and A549 cells were transfected with GFP-Flag (VB200507-2985cmv) or SARS-CoV-2 S-Flag (VB200507-2984jyv) (1.5 μg/ml) constructs using Lipofectamine 3000 reagent (Invitrogen) according to manufacturer’s instructions, and confirmed by observing GFP under fluorescence microscope and western blot analysis of SARS-CoV-2 S and Flag proteins. 48h post transfection, cells were lysed with RIPA lysis buffer containing complete protease inhibitor cocktail and phosphatase inhibitor cocktail (Roche) for the detection of S protein by western blot or ELISA, or resuspended in TRIzol™ reagent (Invitrogen) for the isolation of RNA and subsequent measurement of S mRNA by real-time qPCR.

### In-vitro stimulation of epithelial cells

To examine the effect of SARS-CoV-2 proteins on inflammatory responses in epithelial cells, HEK293T and A549 cells were stimulated with SARS-CoV-2 S1 (RayBiotech, 230-30161), SARS-CoV-2 S2 (Ray Biotech, 230-30163) for 4 hrs. RNA was isolated and measured for the expression inflammatory genes by real-time PCR

### Co-culture of macrophages and epithelial cells

HEK293T-GFP and HEK293T-SARS-CoV-2 S or A549-GFP and A549-SARS-CoV-2 S cells were cultured with THP-1 macrophage-like cells in a ratio of 1:2 (macrophages were twice in number to epithelial cells). Following 16h of co-culture, culture medium was collected, filtered with 0.2 μM filter, and used for ELISA. RNA was isolated and measured for the expression inflammatory genes by real-time PCR.

### Stimulation of macrophages with conditioned medium of S-protein expressed epithelial cells

At 50 – 60 % confluency, HEK293T and A549 cells were transfected with GFP-Flag or SARS-CoV-2 S-Flag (1.5 μg/ml) constructs using Lipofectamine 3000 reagent (Invitrogen) according to manufacturer’s instructions, and confirmed by observing GFP under fluorescence microscope and western blot analysis of SARS-CoV-2 S and Flag. 48h post-transfection, culture medium (conditioned medium) was collected, filtered with 0.2 μM filter and store at −80 °C. At about 85% confluency, culture medium of macrophage like THP-1 cells were replaced with new media containing 30% (V/V) of epithelial cell conditioned medium. After 4 h incubation with conditioned medium, the expression of inflammatory cytokines and chemokines were measured.

### Real-time PCR

Epithelial cells, BMDMs, and THP-1 macrophage-like cells, RAW264.7 cells were lysed in TRIzol™ reagent (Invitrogen). Total RNA was isolated using TRIzol™ reagent (Invitrogen) following the manufacturer’s instructions. Isolated RNA was reverse transcribed into cDNA using iScript (Bio-Rad). Real-time PCR was performed using iTaq Universal SYBR Green Supermix (Bio-Rad). Individual expression data was normalized to GAPDH as described earlier (45). Primers used for qRT-PCR are listed in Table S1.

### In-vitro studies with human peripheral blood mononuclear cells (PBMCs)

Human PBMCs were obtained from STEMCELL™ TECHNOLOGIES (70025), and were cultured in RPMI-1640 medium (R8758, Sigma) supplemented with 10% (v/v) FBS (Sigma) and 1% (v/v) PenStrep (Sigma) and maintained in a 5% CO_2_ incubator at 37°C. After 48 h, hPBMCs were centrifuged and resuspended in fresh medium for 3 h, and then stimulated with SARS-CoV-2 S (500ng/mL) for 4 hrs.

### Inflammatory response of SARS-CoV-2 S protein in mice

WT and *Tlr2^−/−^* mice were intraperitoneally injected with S1 and S2 subunits of SARS-CoV-2 S protein at equal concentration (1ug each/mouse). Blood was collected before S protein administration by cheek puncturing. 15 h following treatment, mice were sacrificed, and blood was drawn from the heart. Serum was separated from the blood by centrifugation and used for the measurement of cytokines by ELISA.

### ELISA

BMDMs and THP-1 macrophage-like cells were lysed in ice-cold RIPA buffer supplemented with complete protease inhibitor and phosphatase inhibitor cocktails (Roche). Protein concentration was measured by the Pierce™ BCA Protein Assay Kit (Thermo Scientific-23227). Serum was isolated from mouse blood by centrifugation at 10,000 RPM for 10 min at 4°C. The concentration of IL-6, IL-1 ß, and TNF-a in cell culture medium and serum was measured using commercially available ELISA kits (R&D Systems). SARS-CoV-2 S protein in cell lysates was detected by SARS-CoV-2 S ELISA kit and following manufacturer’s instruction (RayBiotech, ELV-COVID19S2).

### Western blot

THP-1 macrophage-like cells, BMDM, HEK293T and A549 cells were lysed in ice-cold RIPA lysis buffer containing complete protease inhibitor and phosphatase inhibitor cocktails (Roche), resolved by SDS-PAGE, and transferred onto a PVDF membrane. The membranes were immunoblotted with antibodies against Phospho-NF-κB p65 (3033, Cell Signaling), Phospho-IkBa (9246, Cell Signaling), IkBa (4812, Cell Signaling), Phospho-ERK (4370, Cell Signaling), ERK (4695, Cell Signaling), Phospho-JNK (4668, Cell Signaling), Phospho-AKT (4060, Cell Signaling), AKT (9272, Cell Signaling), Phospho-STAT3 (9145, Cell Signaling), Flag^®^M2 (F1804, Sigma-Aldrich), SARS-CoV-2 S (GTX632605, GeneTex) and ß-actin (A2228, Sigma). Immunoreactive protein bands were detected using ECL super signal west femto substrate reagent (Thermo Scientific).

### Flow cytometric analysis of SARS-CoV-2 overexpressed HEK293T cells

HEK293T cells were transfected with SARS-CoV-2 plasmid. 48 hours following transfection, cells were trypsinized and processed for cell surface staining. Briefly, 0.5×10^6^ cells were resuspended in staining buffer (00-4222-26; eBioscience) and centrifuged at 1500 RPM for 3 min at 4 °C. Cells were then incubated with Fc block CD16/32 monoclonal antibody (14-0161-82, eBioscience) and stained with primary antibody SARS-CoV-2 (GTX632604, GeneTex) for 30 min on ice. After washing with staining buffer once, cells were stained with secondary antibody DyLight 488 (35552, Thermo Scientific) on ice for 1h. Finally, cells were washed twice with staining buffer and acquired in flow cytometer (CytoFLEX-Beckman Coulter). Flow cytometric data were analyzed by FlowJo software.

### Heat inactivation of S2 protein

S2 protein was heated at 95C for 30 in. Native of heat-denatured S2 proteins (500 ng/ml) were used to stimulate THP1 cells and measurement of Cytokines by real-time PCR.

### Statistical Analysis

Data are represented as mean ± SD or SEM. Data were analyzed by Prism8 (GraphPad Software) and statistical significance was determined by two-tailed unpaired Student’s *t* test. *p* < 0.05 was considered statistically significant.

## ACKNOWLEDGEMENT

We would like to thank the UT Southwestern Animal Resource Center (ARC) for maintenance and care of our mouse colony. We thank Dr. Zhijian “James” Chen for sharing *Myd88^−/−^* mice and Dr. Esra Akbay for sharing A549 cells. Hasan Zaki is supported by The National Institute of Diabetes and Digestive and Kidney Diseases (NIDDK) of the National Institute of Health (NIH) under Award Number R01DK125352 and Cancer Prevention and Research Institute of Texas (CPRIT) Individual Investigator Award (RP200184). Rashmin C. Savani holds the William Buchanan Chair in Pediatrics and is funded by a Sponsored Research Agreement with Mallinckrodt Pharmaceuticals, Inc for an unrelated project.

## Author contributions

S.K. designed and performed experiments, and analyzed data, and helped in manuscript writing. M.S.S. and C.L. helped in experiment. R.C.S. provided *Tlr2^−/−^* mice. J.S. and R.C.S. helped in experimental design, and reviewed the manuscript. H.Z. conceived the study, designed experiments, and wrote the manuscript.

