## Supplemental Figures for "SARS-CoV-2 spike protein induces inflammation via TLR2-dependent activation of the NF-κB pathway"

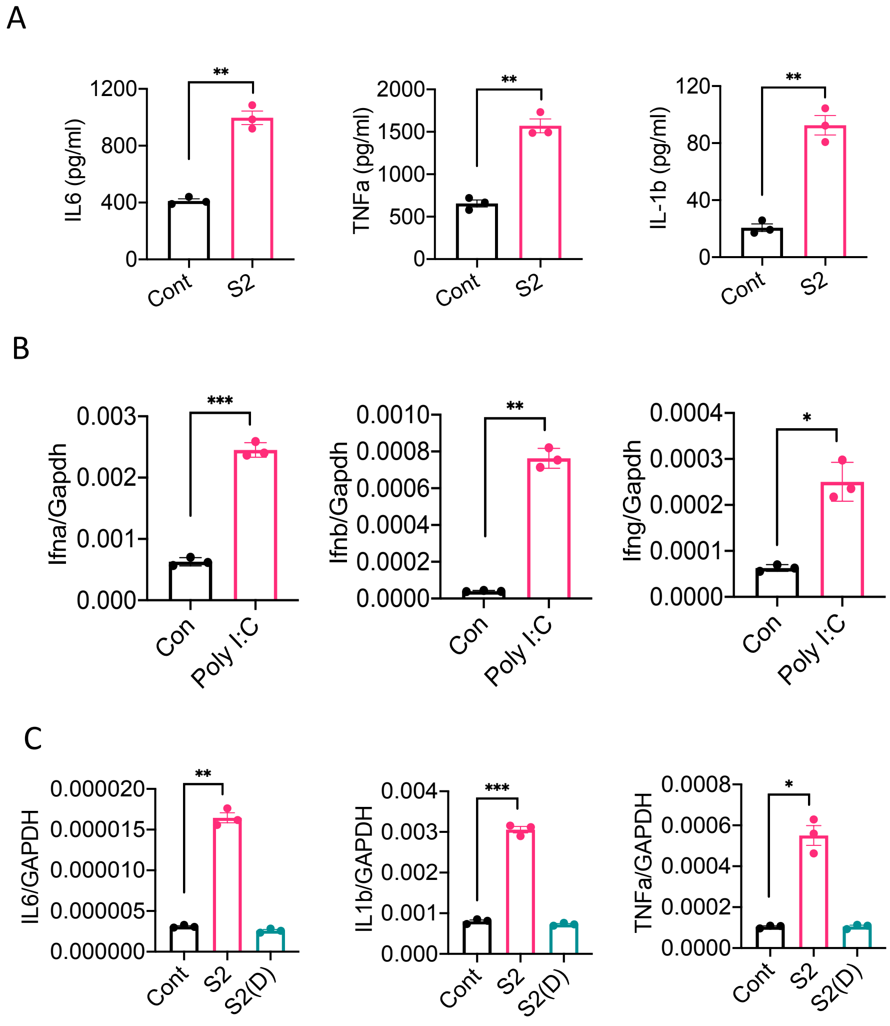


**Figure S1. SARS-CoV-2 S protein induces inflammatory cytokines in macrophages.** (A) THP-1 cells were stimulated with S2 protein. 4h post stimulation, culture supernatants were collected, centrifuged, and analyzed for IL-6, IL-1β, and TNFα by ELISA. (B) THP-1 cells were stimulated with PolyI:C (1μg/ml). The expression of interferons was measured by real-time PCR. (C) THP1 cells were stimulated with native S2 or heat-denatured S2 (500 ng/ml). The induction of IL6, IL1b, and TNFa was measured by real-time PCR at 4h post stimulation. Data represent mean ± SD (n=3); **$p < 0.001$ by unpaired Student's $t$ test. Experiments were repeated two times and data of a representative experiment is presented.

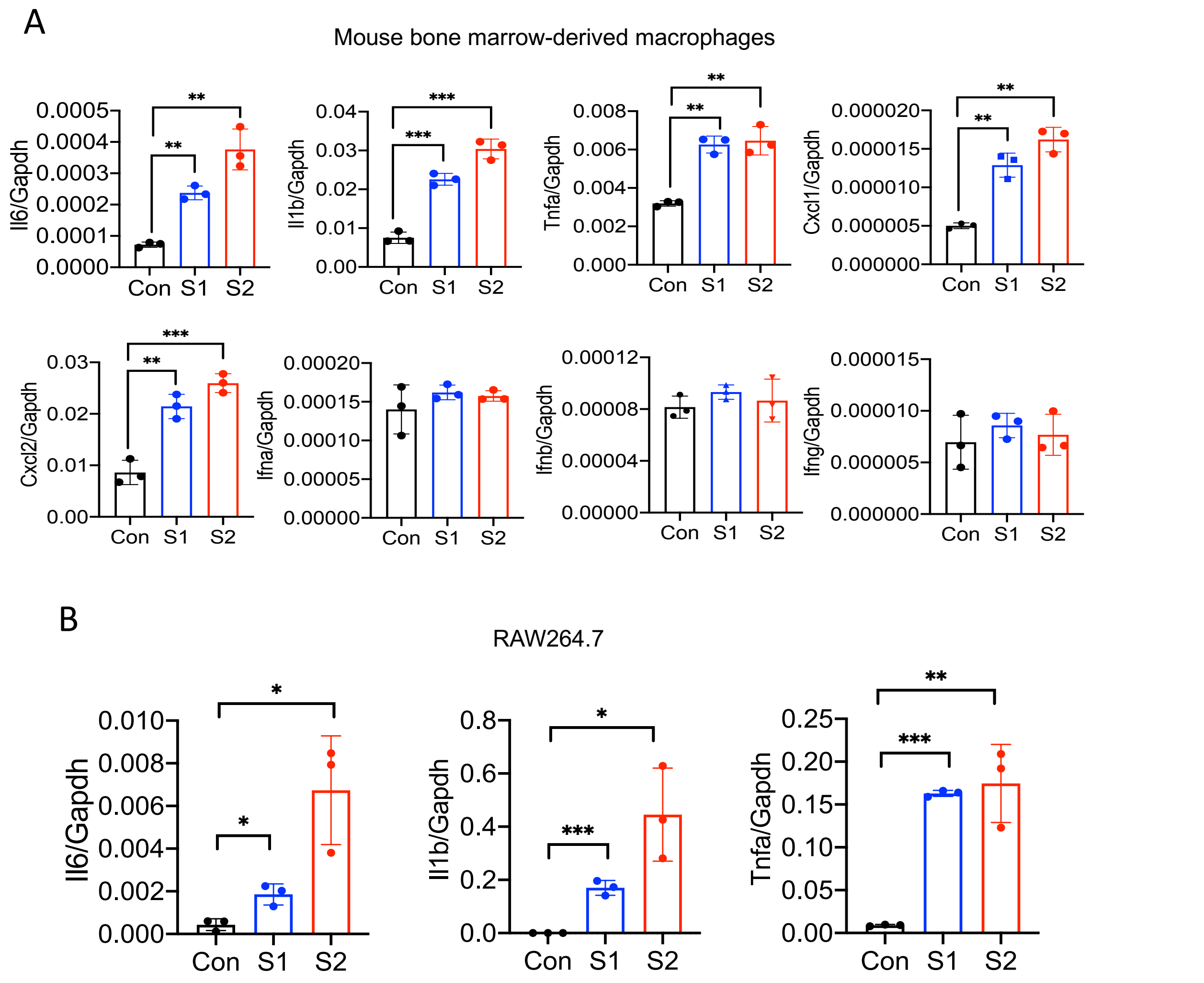


**Figure S2. Mouse macrophages are stimulated by SARS-CoV-2 S protein.** (A) Bone marrow-derived macrophages from WT mice were stimulated with S1 and S2 proteins (500 ng/ml) for 4 h. The expression of *Il6*, *Il1b*, *Tnfa*, *Cxcl1*, *Cxcl2, Ifna, Ifnb, Ifng* was measured by real-time PCR. (B) RAW264.7 murine macrophage cells were stimulated with S1 or S2 (500 ng/ml) for 4h. The expression of *Il6, Il1b*, and *Tnfa* was measured by real-time PCR. Data represent mean ± SD (n=3); *$p < 0.05$, **$p < 0.001$, ***$p < 0.0001$ by unpaired Student's *t* test. Experiments were repeated two times and data of a representative experiment is presented.

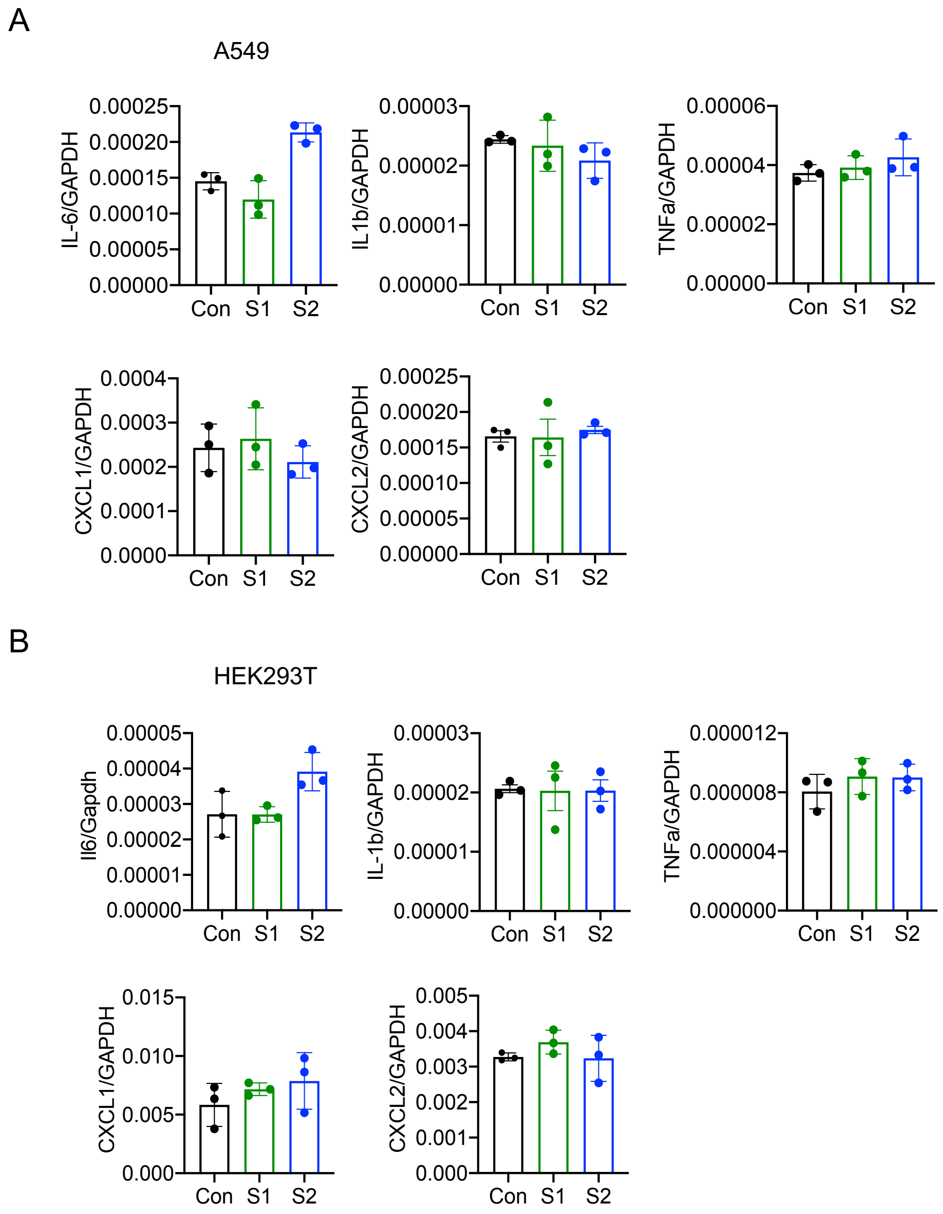


**Figure S3. Epithelial cells do not respond to SARS-CoV-2 S protein acutely.** (A-B) A549 or HEK293T cells were incubated with SARS-CoV-2 S1 or S2 (500 ng/ml) proteins for 4 h. The expression of inflammatory cytokines and chemokines was measured by real-time PCR. Data represent mean ± SD (n=3). Experiments were repeated two times and data of a representative experiment is presented.

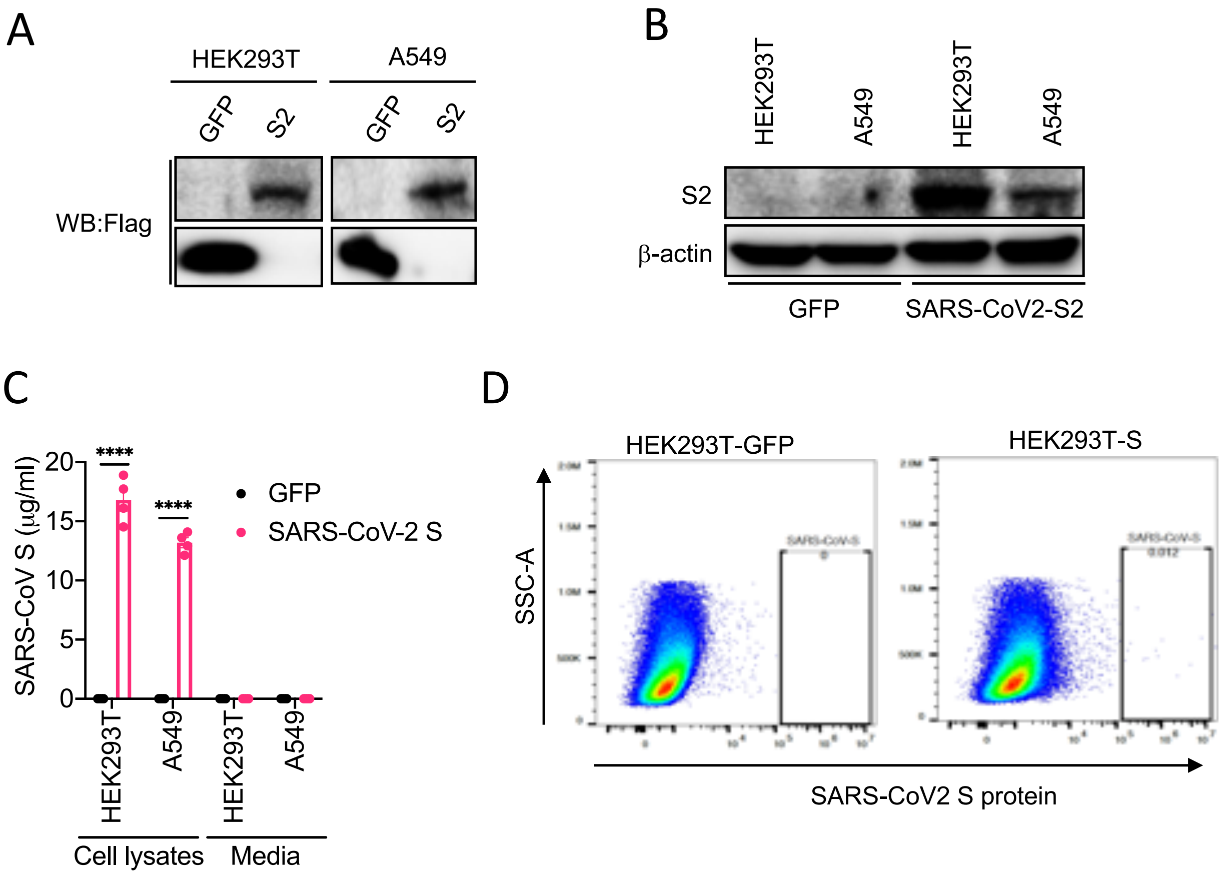


**Figure S4. Expression analysis of SARS-CoV-2 S protein in cultured cells.** HEK293T or A549 cells were transfected with plasmids containing flag-tagged S or green fluorescent protein (GFP). (A-B) 48h post transfection, cell lysates were collected and the expression of S was measured by western blot analysis of Flag (A) and S (B) using respective antibodies. (C) Cell culture supernatants and cell lysates were collected at 48h after transfection, and analyzed for S protein by ELISA. (D) The expression of S on the cell surface of HEK293T-S cells was measured by flow cytometry following surface staining of S proteins. Data represent mean ± SD; ****$p < 0.0001$ by unpaired Student's $t$ test. Experiments were repeated two times and data of a representative experiment is presented.

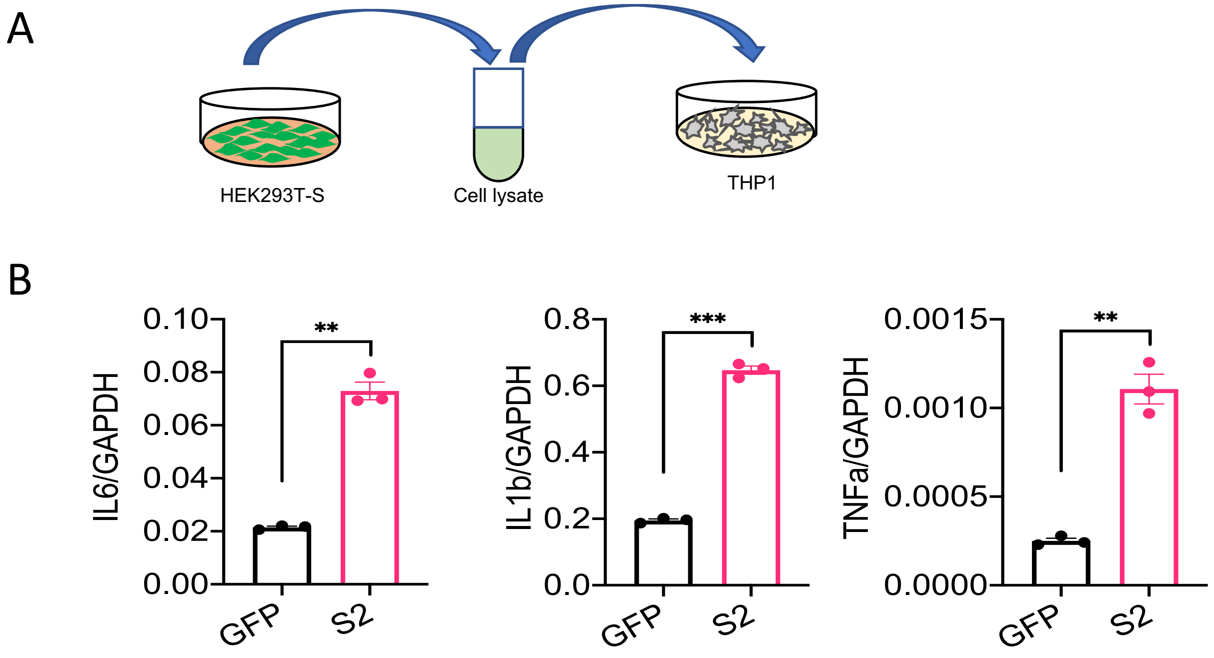


**Figure S5. Cell lysate of HEK293T cells expressing S protein activates macrophages and induced inflammatory cytokines.** (A) S or GFP (control) proteins were overexpressed in HEK293T cells. Cells were sonicated, and cell lysate supernatants were collected. THP-1 cells were incubated with these cell lysate supernatants for 4 h. (B) The expression of *IL6, IL1b,* and *TNFa* in THP cells was measured by real-time qPCR. Data represent mean ± SD (n=3); $^{**}p < 0.001$, $^{***}p < 0.0001$ by unpaired Student's *t* test. Experiments were repeated two times and data of a representative experiment is presented.

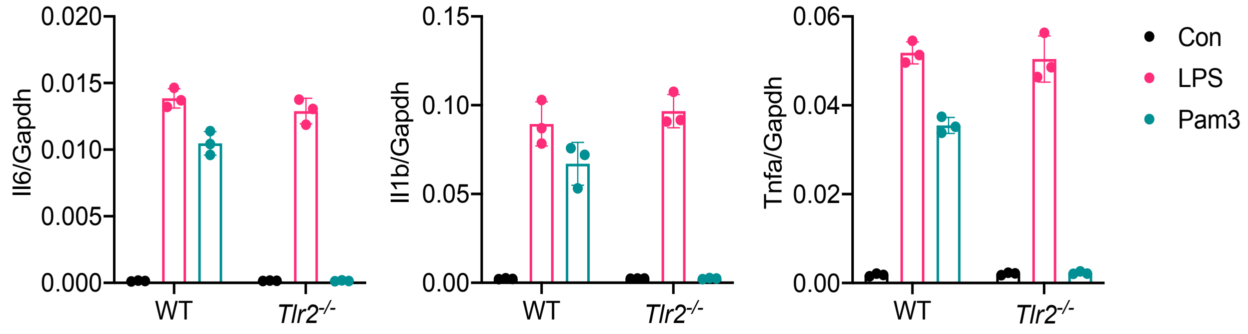


**Figure S6. Macrophages of *Tlr2*$^{-/-}$ mice are defective in sensing TLR2 ligand Pam3CSK4.** BMDMs from WT and *Tlr2*$^{-/-}$ mice were stimulated with LPS (1μg/l) or Pam3CSK4 (1μg/ml) for 4 h. The expression of inflammatory cytokines was measured by real-time PCR. Data represent mean ± SD (n=3).
